# Competence induction of homologous recombination genes protects pneumococcal cells from genotoxic stress

**DOI:** 10.1101/2024.07.12.603235

**Authors:** David De Lemos, Anne-Lise Soulet, Violette Morales, Mathieu Berge, Patrice Polard, Calum Johnston

## Abstract

Homologous recombination (HR) is a universally conserved mechanism of DNA strand exchange between homologous sequences, driven in bacteria by the universal recombinase RecA. HR is key for the maintenance of bacterial genomes via replication fork restart and DNA repair, as well as for their plasticity via the widespread mechanism of natural transformation. Transformation involves the capture and internalisation of exogenous DNA in the form of single strands (ssDNA), followed by chromosomal integration via HR. In the human pathogen *Streptococcus pneumoniae*, transformation occurs during a transient, stress-induced physiological state called competence. RecA and its partner DNA branch migration translocase RadA both cooperate in HR during transformation and in some recombinational DNA repair pathways of genome maintenance. Both *recA* and *radA* genes are basally expressed and transcriptionally induced during competence. In this study, we explored the importance of competence induction of *recA* and *radA* expression in transformation and genome maintenance processes. We confirmed that competence induction of *recA* was important for optimal transformation, but found this was not the case for *radA*. In contrast, the competence induction of both genes was required for optimal tolerance faced with transient exposure to the lethal genotoxic agent methyl methanesulfonate (MMS). However, this was not the case for another DNA-damaging agent, norfloxacin. These results show that competence induction of HR effectors is important for the increased tolerance to genotoxic stress provided to competent pneumococcal cells. This reinforces the finding that pneumococcal competence is a stress-sensing mechanism, transiently increasing the expression of some genes not to optimise transformation but to improve survival faced with specific lethal stresses.

**Importance:** Homologous recombination (HR) is a mechanism of DNA strand exchange important for both the maintenance and plasticity of bacterial genomes. Bacterial HR is driven by the recombinase RecA along with many accessory partner proteins, which define multiple dedicated pathways crucial to genome biology. Thus, a main mechanism of genome plasticity in bacteria is natural genetic transformation, which involves uptake and chromosomal integration of exogenous DNA via HR. In the human pathogen *Streptococcus pneumoniae*, transformation occurs during a transient, stress-induced physiological state called competence. RecA and the helicase RadA are key for both genome maintenance and transformation, and both are over-produced during competence. Here, we explore the importance of this over-production for transformation and genome maintenance, quantified by tolerance to genotoxic stress. While over-production of RecA was important for both processes, over-production of RadA was required only for genotoxic stress tolerance. This highlights the importance of competence as a stress-responsive mechanism, with induction of HR genes important for genotoxic stress tolerance.

## Introduction

Homologous recombination (HR) is central to the maintenance and plasticity of bacterial genomes. HR is a universally conserved mechanism of exchange between homologous DNA strands that is driven by the universal homologous recombinase RecA in bacteria^1^. HR plays key roles in several vital pathways of genome maintenance, including replication fork restart^2^ and repair of single stranded DNA (ssDNA) gaps or double-stranded DNA (dsDNA) breaks^3^. In addition, HR is essential for the widespread bacterial horizontal gene transfer mechanism of natural transformation^4^. In all of these processes, HR begins with the loading of RecA onto nascent ssDNA by dedicated recombination mediator proteins (RMPs)^5^, which promote ATP-dependent nucleofilamentation of RecA to generate a dynamic polymer known as the presynaptic HR filament. This intermediate then undergoes a homology search within the chromosome and, once a complementary sequence is found, directs ssDNA exchange to form a displacement loop (D-loop). The D-loop is then processed and resolved to restore the DNA duplex, leading to replication fork restart, repair of damaged DNA or integration of new genetic content by transformation (Figure 1A).

**Figure 1:**
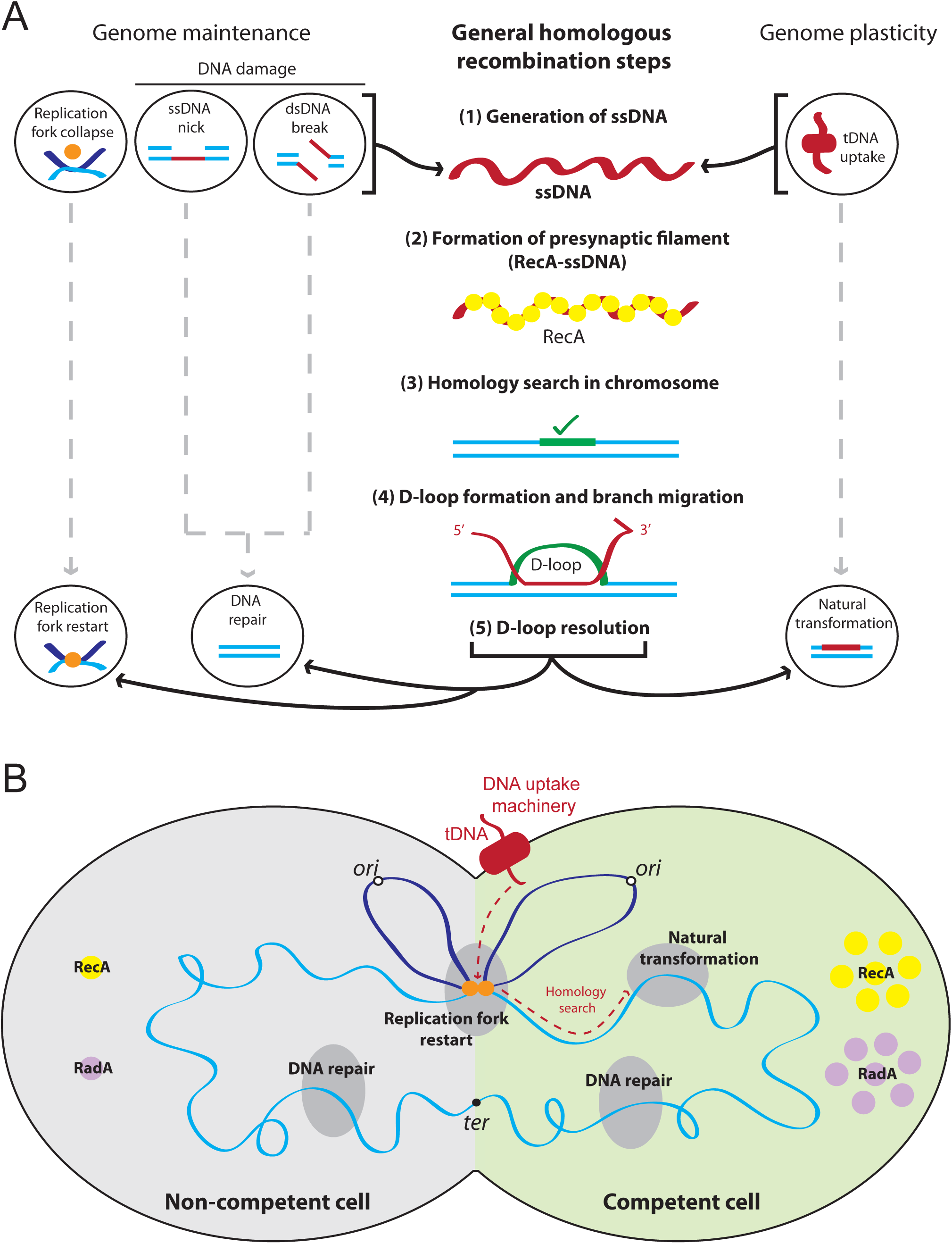
Homologous recombination (HR) pathways in competent and non-competent pneumococcal cells. (A) General steps conserved among HR pathways in the pneumococcus. (1) The first step of HR involves the generation of ssDNA within the cell, which can occur via replication fork collapse, DNA damage (ssDNA nicks or dsDNA breaks) or uptake of ssDNA during the genome plasticity mechanism of natural transformation. (2) The homologous recombinase RecA is the loaded onto the ssDNA, and polymerises to form a presynaptic filament. (3) RecA mediates homology search within the chromosome. (4) Strand exchange between homologous sequences forms a recombination intermediate known as a D-loop, with branch migration by helicases including RadA extending this structure. (5) Further processing and D-loop resolution results in replication fork restart, DNA repair or integration of transforming ssDNA, respectively. Light blue line, chromosomal DNA; dark blue line, neosynthesized chromosomal DNA; orange dot, replisome; dark red lines, ssDNA; dark red oblong, transformation machinery; yellow circles, RecA molecules; green line, homologous DNA in chromosome. (B) HR pathways in competent and non-competent pneumococcal cells and implication of both RecA and RadA. In non-competent cells, HR is involved in genome maintenance pathways including replication fork restart and DNA repair. In addition, both RecA and RadA are expressed at basal levels. In competent cells, HR is still involved in replication fork restart and DNA repair, but also in transformation, where tDNA is internalised at midcell, and accesses the chromosome via active replication forks, from which homology search may emanate^54,55^. Once homology is found, HR promotes integration of tDNA into the recipient chromosome. Both *recA* and *radA* are induced during competence, resulting in increased cellular levels of RecA and RadA in competent cells. Grey half-cell, non-competent cell; green half-cell, competent cell; dark grey ovals, HR pathways; light blue line, chromosomal DNA; dark blue line, neosynthesized chromosomal DNA; orange dot, replisome; dark red line, tDNA; dark red oblong, transformation machinery; yellow circles, RecA molecules; purple circles, RadA molecules; white dots, origins of replication; black dot, terminus of replication; red dotted line, path of tDNA from internalisation to integration.

Natural transformation involves the capture and internalization of exogenous DNA into the cell cytoplasm in the form of single strands, which are then integrated into the recipient chromosome by HR^4^. This promotes the acquisition of new genetic traits, notably mediating the spread of antibiotic resistance and vaccine escape. In the human pathogen *Streptococcus pneumoniae* (the pneumococcus), transformation occurs during a transient physiological state called competence, which is a tightly regulated genetic program that lasts ∼20 minutes at the cellular level^6^. During pneumococcal competence, around 100 genes are induced, including genes encoding the multiprotein transformation machinery^7–9^. Pneumococcal competence is induced in response to environmental factors such as antibiotic or genotoxic stress^10–13^, and has been proposed as an analogue to the SOS system, which is absent in this species^14^. In addition, pneumococcal competence modulates the survival of cells exposed to certain stresses including antibiotics and genotoxic agents^15,16^.

Pneumococcal competence induction occurs in two transcriptional waves, with early competence genes induced by the response regulator ComE upon its phosphorylation by the membrane-bound ComD kinase^6^. Late competence genes are induced by an alternative sigma factor σ^X^, itself induced by phosphorylated ComE. Twenty-two of the >100 genes induced during competence are directly involved in transformation, and these are all controlled by σ^X17^. Although the majority of these are not expressed outside of competence, a few exceptions exist. These include the HR genes *recA* and *radA*, both of which encode proteins central to genome maintenance and transformation^18,19^. These genes are expressed at a basal level as well as being induced during competence^7–9^. Pneumococcal cells lacking *recA* showed increased sensitivity to multiple types of DNA damages such as UV exposure or the DNA- damaging agent methyl methanesulfonate (MMS), and displayed almost no transformation activity^12,18,20^. RadA is involved in branch migration of three-strand DNA molecules produced by RecA in *Escherichia coli*^21^. Pneumococcal RadA was revealed to be a hexameric DnaB-like helicase, extending ssDNA incorporation at D-loops during transformation^19,22^. In addition, pneumococcal RadA is important for DNA damage, since cells lacking *radA* were more sensitive to MMS^19^.

In non-competent pneumococcal cells, the basal expression of *recA* and *radA* provides the growing cells with a pool of RecA and RadA molecules to face replication fork restart and DNA repair events occurring on the genome (Figure 1B). In competent cells, it is unclear if σ^X^ induction of both *recA* and *radA* expression only serves transformation or also genome maintenance concurrently (Figure 1B). So far, only the importance of *recA* induction for transformation has been established^23^. We recently established that competence itself increases tolerance to genotoxic stress, independent of transformation^16^. However, whether induction of *recA* and/or *radA* are important for this tolerance is unknown. Here, we explored the importance of the competence induction of pneumococcal *recA* and *radA* for their key roles in genome maintenance or transformation. Unexpectedly, unlike *recA*, competence induction of *radA* was not required to ensure optimal transformation. In contrast, we found that the competence induction of both *recA* and *radA* was important for competence-mediated tolerance to the genotoxic agent MMS. Competence is induced in response to cellular stresses including DNA damage^10–12^, and historically has been seen as a conduit for transformation. However, recent studies have linked competence with other important cellular processes such as biofilm formation^24,25^, virulence^26–29^, and transmission^30^ and to foster multilevel heterogeneity within a clonal population^16^. Competence is thus a general stress response of which transformation is one of multiple facets. One facet of this stress response is to increase the ability of cells to tolerate genotoxic stress, and here we show that the competence induction of key HR mediators RecA and RadA is central to this ability.

## Results

### σ^X^ *cin* box mutations prevent competence induction of the *recA* and *radA* operons

Pneumococcal *recA* and *radA* genes are included in complex operons. The *recA* gene is downstream of the *cinA* gene and upstream of the *dinF*^31^ and *lytA*^32^ genes (Figure 2A). Expression of *recA* is controlled by a σ^X^-induced promoter upstream of *cinA*^7–9^ and a constitutive promoter upstream of *recA*. The *radA* gene is part of a three gene operon also containing the dUTP pyrophosphatase gene *dut*^9^ and a gene of unknown function (*spr0024*), controlled by a constitutive promoter and a σ^X^-specific promoter (Figure 2B). A downstream operon consisting of genes encoding the essential pneumococcal carbonic anhydrase Pca^33^ and the degenerate protease PrsW^34^ is also controlled by the same competence promoter. No other proteins produced by these two operons are involved in transformation, although a link between CinA and transformation has been suggested^35^.

**Figure 2:**
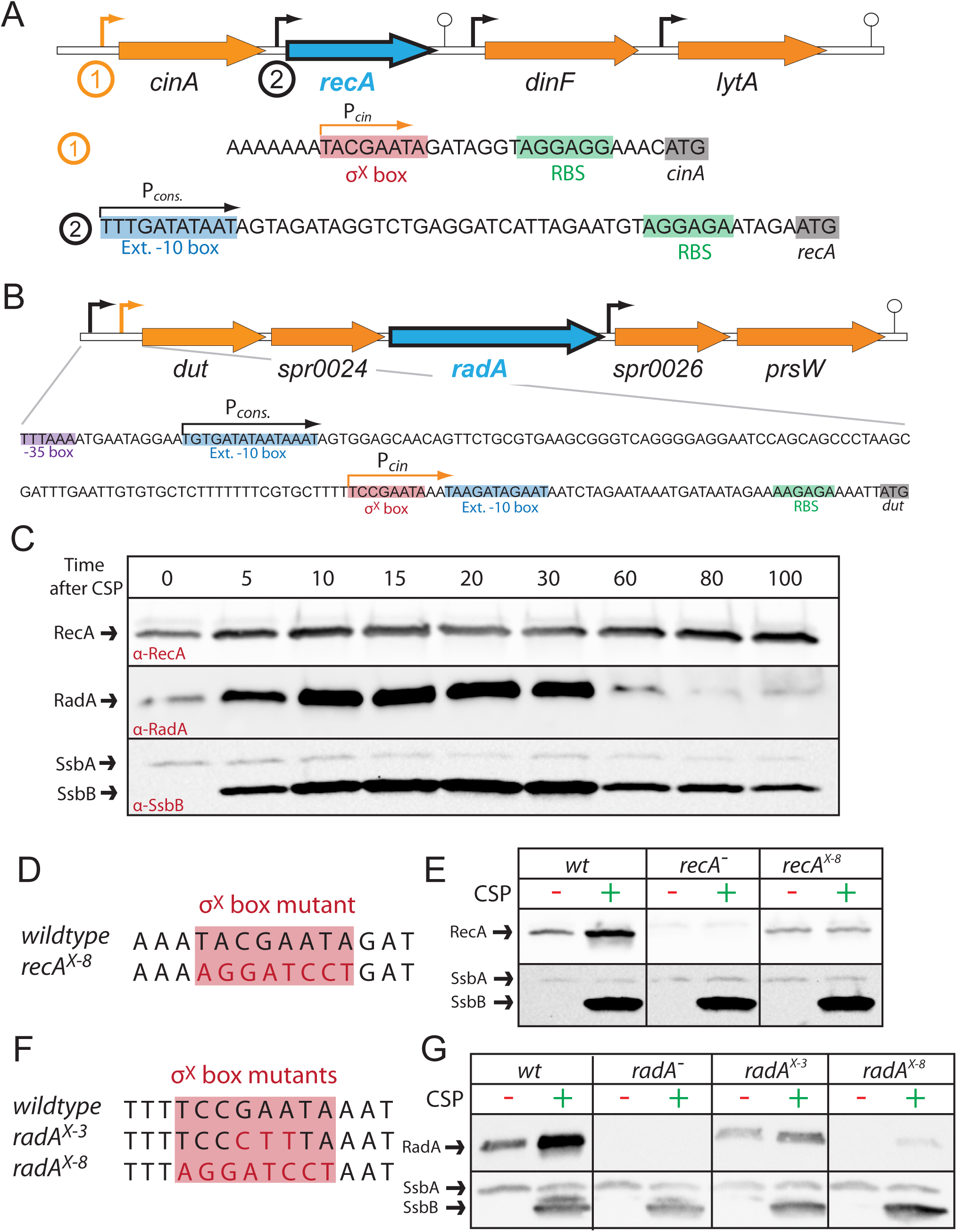
Genetic context, expression and σ^X^ box mutants of *recA* and *radA* operons. (A) Genetic context of the pneumococcal *recA* locus. The *recA* gene is controlled by two promoters, a constitutive promoter (black arrow) and a competence-inducible promoter upstream of *cinA* (orange arrow). Downstream, *dinF* and *lytA* are controlled by their own promoters (black arrows), but also induced via the *cinA* competence promoter^7–9^. Black arrows, constitutive promoter (P*_cons_.*); orange arrow, P*_cin_* promoter; stalked circle, terminator; grey bases, start codon; red bases, σ^X^ box; green bases, ribosome-binding site (RBS); blue bases, extended -10 box. Arrows indicate promoter orientation. (B) Genetic context of the pneumococcal *radA* locus. The *radA* gene is in an operon which, with *dut* and *spr0024*, which is controlled by two promoters, a constitutive promoter (black arrow) and a competence promoter (orange arrow). The downstream operon consisting of *psrW* and *spr0026*, controlled by a second constitutive promoter, is also induced during competence via the same promoter. Black arrows, constitutive promoter (P*_cons._*); orange arrow, P*_cin_* promoter; stalked circle, terminator; grey bases, start codon; red bases, σ^X^ box; green bases, ribosome-binding site (RBS); purple bases, -35 box; blue bases, extended -10 box. Arrows indicate promoter orientation. (C) Kinetics of RecA and RadA expression before, during and after competence. Western blot using α-RecA, α-RadA and α-SsbB antibodies to follow protein levels in cells prior to and after exposure to CSP to induce competence. SsbB antibodies recognize both SsbB and SsbA. (D) Alignment of wildtype *recA* operon σ^X^ box and mutated σ^X^ box *x-8*. Red highlight, σ^X^ box; red bases, mutated bases compared to wildtype. (E) Cellular levels of RecA in competent and non-competent *x-8* σ^X^ box mutant strains. Western blot using α-RecA and α- SsbB antibodies on cellular extracts 15 minutes after addition (or not) of CSP to induce competence. (F) Alignment of wildtype *radA* operon σ^X^ box and mutated σ^X^ boxes *x-3* and *x-8*. Red highlight, σ^X^ box; red bases, mutated bases compared to wildtype. (G) Cellular levels of RadA in competent and non-competent *x-3* and *x-8* σ^X^ box mutant strains. Western blot using α-RadA and α-SsbB antibodies on cellular extracts 15 minutes after addition (or not) of CSP to induce competence.

The fact that both *recA* and *radA* are controlled by two promoters, one constitutive and one competence-inducible, suggests that basal expression of these genes is sufficient to ensure genome maintenance in non-competent cells, but that a boost in cellular levels is required to ensure transformation in competent cells and possibly genome maintenance during this transient differentiation state. The induction of *recA* and *radA* during competence is dependent on the alternative sigma factor σ^X^, which recognises an 8 bp sequence (σ^X^ box) in the promoters^8,36^. Comparing the production of RecA and RadA to the competence-specific protein SsbB^37,38^ and its basally expressed paralogue SsbA^39^ after competence induction showed that, as expected, RecA, RadA and SsbA were present prior to CSP addition, unlike SsbB (Figure 2C). Once competence was induced, expression of *recA, radA* and *ssbB* increased, reaching a peak 20 minutes after CSP addition. Once competence was shut-off, cellular levels of RadA and SsbB decreased as cells resumed growth, while RecA remained present at higher levels in post-competent cells. This may be because the basal level of expression of *recA* is higher than *radA*, or that RecA is more stable post-competence than RadA or SsbB, as has been reported previously for certain other competence proteins^40^. To explore the importance of competence induction of HR genes *recA* and *radA* in transformation and genome maintenance, we generated strains with σ^X^ box mutations designed to abrogate the competence induction of the *recA* and *radA* genes. Replacing the eight bases of the *recA* σ^X^ box (*recA^X-^*^8^; Figure 2D) resulted in loss of competence induction without altering basal expression (Figure 2E), and this strain was compared with wildtype (wt) and *recA^-^* strains in this study. In contrast, replacing the eight bases of the *radA* σ^X^ *cin* box (*radA^X-^*^8^; Figure 2F) strongly affected competence induction but also affected basal expression (Figure 2G). This may be due to the proximity of the basal and competence promoters of *radA*, along with the fact that the basal promoter is upstream of the competence promoter (Figure 2B), unlike for *recA* (Figure 2A). Replacing three central bases of the *radA* σ^X^ box (*radA^X-^*^3^; Figure 2F) reduced *radA* expression to a lesser extent than *radA^X-^*^8^, with again minor competence induction observed (Figure 2G). As a result, since neither strain represented a perfect abrogation of *radA* competence induction, both strains were studied in parallel with wt and *radA^-^* strains in this study. Complementation strains with *recA* or *radA* expressed ectopically from the IPTG- inducible P*_lac_* promoter^26^ at the CEP (CEP*_lac_-recA*, CEP*_lac_-radA*) expression platform^41^ were generated (Figure S1) and used when appropriate to complement competence induction mutants.

### Competence induction of the *radA* operon is not required for optimal transformation

Two main types of transformation events exist, integration of a single nucleotide polymorphism (SNP) and integration of a heterologous gene cassette (HGC), both flanked by homologous DNA (Figure 3A). The competence induction of *recA* was previously shown to be important for transformation using a strain where the competence promoter was inactivated by insertion of an antibiotic resistance cassette^23^. To confirm this, we compared transformation of an *rpsL41* point mutation centrally located on a ∼3,5 kb PCR fragment (*rpsL41c*^22^; Figure 3B) at saturating and non-saturating conditions in wt, *recA^-^*, *recA^X-^*^8^ and complemented *recA^X-^*^8^, CEP*_lac_-recA* strains. At high tDNA concentrations (2,500 pg µL^-1^, referred to as saturating tDNA conditions hereafter), wt cells transformed at almost 100 %, while only residual transformants were observed in *recA^-^*cells (Figure 3C), as previously reported^18^. The *recA^X-8^*strain transformed ∼15-fold less than wildtype (Figure 3C), showing that competence induction of *recA* was important for transformation. Complementation of this *recA^X-8^*mutation via CEP*_lac_-recA* reduced the deficit to ∼4-fold compared to wt, a partial complementation which may be explained by the fact that maximal expression levels of CEP*_lac_-recA* do not match those of *recA* during competence (Figure S1A), or by a role in transformation for the competence induction of another operon member. Reducing the concentration of tDNA reduced the overall levels of transformation, confirming that the tDNA was non-saturating (25 and 2,5 pg µL^-1^), but similar differences in transformation efficiency were observed compared to wildtype at all concentrations tested (Figure 3C), confirming a general importance of *recA* competence induction for transformation.

**Figure 3:**
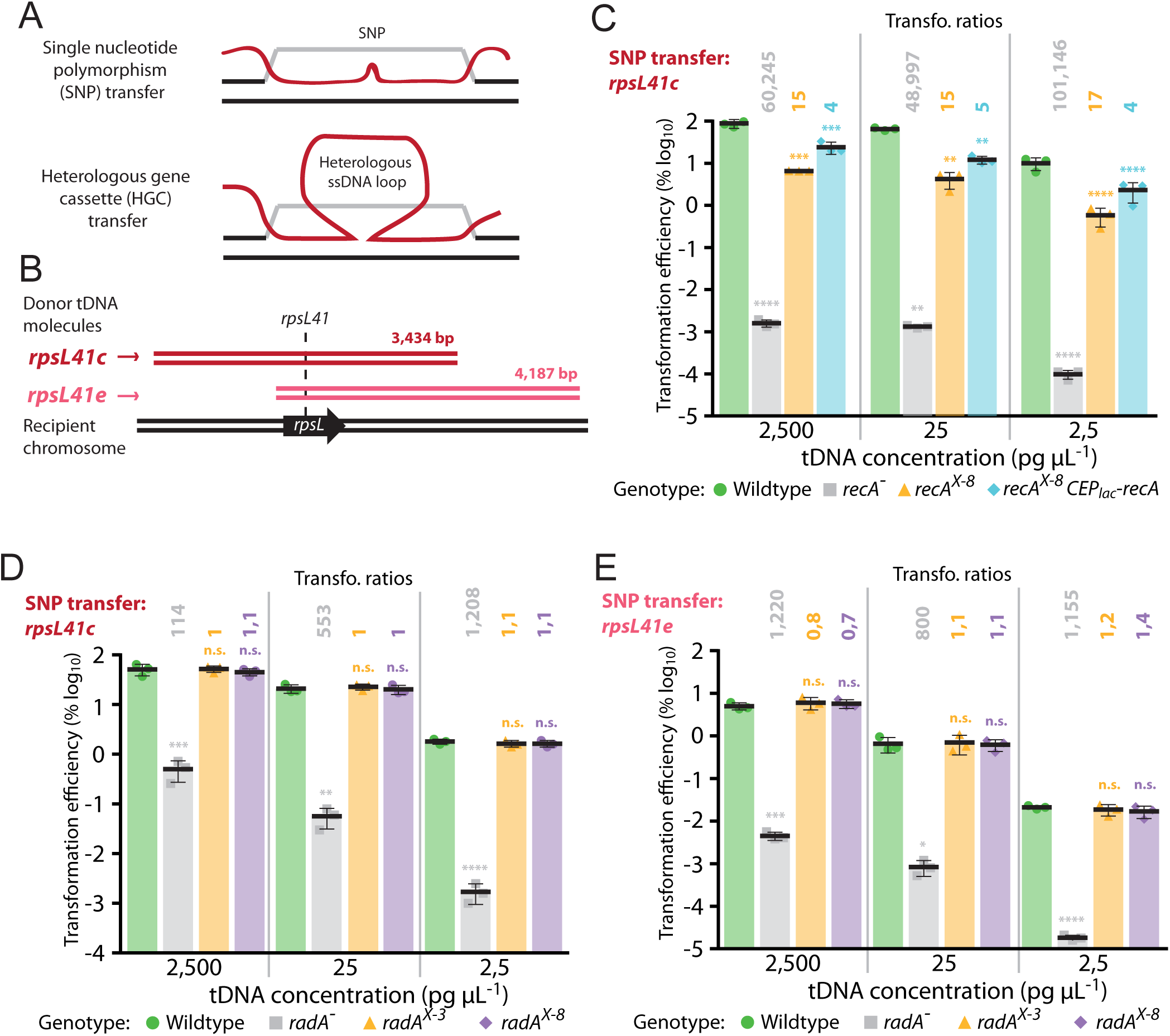
Competence induction of *recA* but not *radA* is important for transformation. (A) Schematic representation of chromosomal integration of single nucleotide polymorphism (SNP) and heterologous gene cassette (HGC) by transformation. These structures are resolved by replication into one wildtype and one transformed daughter chromosome. For HGC integration, the heterologous ssDNA loop represents the heterologous sequence, which is integrated into a daughter chromosome by replication. (B) Donor DNA fragments and recipient chromosome identity. Donor fragments possess the *rpsL41* point mutation, which provides streptomycin resistance (Sm^R^). In *rpsL41c* (dark red), the *rpsL41* point mutation is located centrally while in *rpsL41e* (light red), it is present at the 5’ extremity of the DNA fragment. (C) Competence induction of *recA* is important for transformation in both saturating (2,500 pg mL^-1^) and non-saturating (25 and 2,5 pg mL^-1^) tDNA conditions. Transformation efficiency of *rpsL41c* DNA fragments into wildtype (R1501, green), *recA^-^* (R4857, gray), *recA^X-8^* (R5077, orange) and *recA^X-8^,* CEP*_lac_-recA* (R5077, blue) strains. Error bars representative of triplicate repeats, with each data point shown. Transformation ratios calculated by comparing average transformation efficiencies of test strains with wildtype. Statistical analyses tested significant difference between wildtype and test strains. ****, p < 0,001; ***, p > 0,005; **, p > 0,01. (D) Competence induction of *radA* is not necessary for optimal transformation of a point mutation central on the tDNA molecule (*rpsL41c*). tDNA in saturating (2,500 pg mL^-1^) and non-saturating (25 and 2,5 pg mL^-1^) conditions. Strains tested; wildtype (R1501, green), *radA^-^* (R2091, gray), *radA^X-3^* (R4091, orange) and *radA^X-8^* (R4635, purple). Error bars representative of triplicate repeats, with each data point shown. Transformation ratios calculated and statistical analyses carried out as in panel C. Statistical analyses tested significant difference between wildtype and test strains. ****, p < 0,001; ***, p < 0,005; **, p < 0,01, n.s., p > 0,05. (E) Competence induction of *radA* is not necessary for optimal transformation of a point mutation eccentric on the tDNA molecule (*rpsL41e*). tDNA concentrations, strains and colour schemes as in panel D. Transformation ratios calculated and statistical analyses carried out as in panel C. ****, p < 0,001; ***, p < 0,005; *, p < 0,05, n.s., p > 0,05.

Since *radA* is specifically induced during competence^7–9^, and key for transformation^19,22^, we hypothesized that the induction of *radA* could be similarly important for transformation. To explore this hypothesis, we repeated the transformation assay with wt, *radA^-^*, *radA^X-3^* and *radA^X-8^* strains transformed with the ∼3,5 kb PCR fragment *rpsL41c* as tDNA. When saturating DNA conditions were used (2,500 pg µL^-1^), no difference in transformation efficiency was observed for either the *radA^X-3^* or the *radA^X-8^*strains compared to wt (Figure 3D). The control *radA^-^*strain showed a 2-log deficit in transformation efficiency compared to wildtype (Figure 3D), in line with previously reported results^22^. Competence induction of the *radA* operon remained unnecessary for optimal transformation when non-saturating tDNA concentrations were used (25 and 2,5 pg µL^-1^, Figure 3D). RadA was previously shown to be important for extension of transformation D-loops, revealed by an increased importance for transformation of a point mutation on the extremity of a PCR fragment (*rpsL41e*, eccentric, Figure 3B)^22^. Repeating the experiment with this tDNA donor molecule revealed that competence induction of *radA* was not required for optimal transformation of eccentric point mutations either, irrespective of the tDNA concentration used (Figure 3E). In addition, transforming cells with saturating (2,500 pg µL^-1^) or non-saturating (25 and 2,5 pg µL^-1^) concentration of a tDNA harbouring a heterologous cassette providing kanamycin resistance (Figure 4A) showed that induction of the *radA* operon was not required for efficient HGC transformation (Figure 4B). Transforming with PCR products targets the same chromosomal location and thus theoretically generates a maximum of two concurrent D-loops on a replicating pneumococcal chromosome. However, transformation with otherwise homologous genomic DNA possessing a seletable mutation should generate many D-loops on the chromosome within a single transforming cell, which may titrate RadA and render its induction during competence important for optimal transformation. To test this, the same strains, along with two other strains complementing the lack of competence induction by ectopic, IPTG-inducible expression of *radA* from the P*_lac_* promoter^42^ (Figure S1B), were transformed with homologous genomic DNA containing the *rpsL41c* point mutation. Transformations were carried out in saturating (2,500 pg µL^-1^) and non-saturating (25 and 2,5 pg µL^-1^) concentrations of tDNA. No differences in transformation efficiency compared to wt were observed, except for the *radA^X-8^* strain, which showed a statistically significant 2-fold decrease in saturating tDNA conditions (Figure 4C). This could suggest that in these conditions, competence induction is required for optimal transformation, but since basal *radA* expression is lower in this strain (Figure 2G), and no difference is observed for *radA^X-3^*, it is likely that a correct basal level of *radA* expression would ensure optimal transformation in these conditions. Collectively, these results suggest that the induction of *radA*, or indeed the other four genes induced by the same competence promoter, was not required for optimal transformation efficiency, even for transformation contexts in which RadA was particularly important^22^ or titrated by multiple D-loops. In conclusion, competence induction of *recA* is important for transformation, while competence induction of *radA* is not.

**Figure 4:**
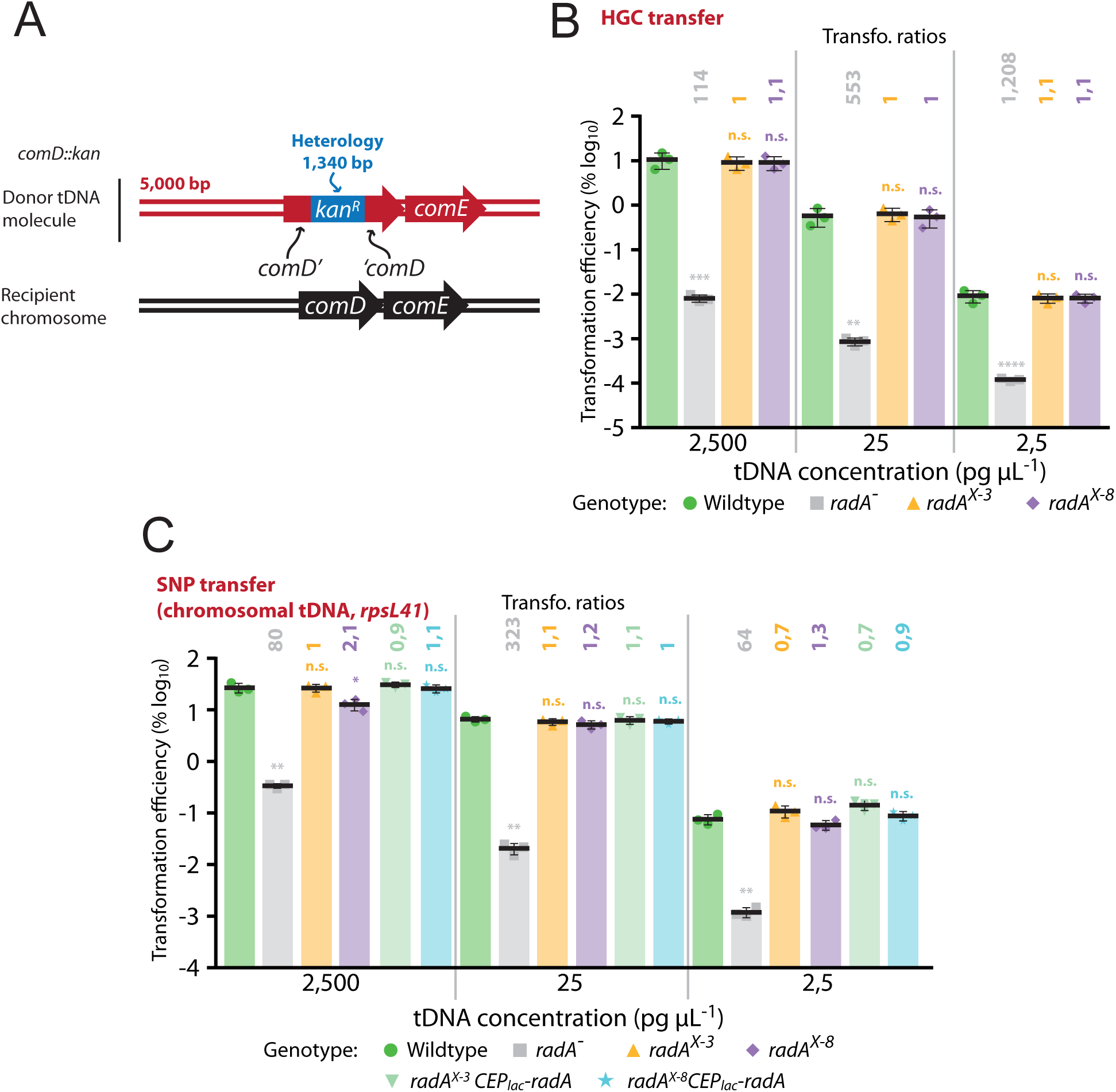
Competence induction of *radA* operon is not required for integration of heterologous cassettes by transformation. (A) Donor DNA fragments and recipient chromosome identity for HGC transfer. Transformation inserts a kanamycin resistance cassette into the *comD* gene, integrating 1,340 bases of heterology. (B) Competence induction of *radA* is not necessary for optimal transformation of heterologous cassettes. tDNA concentrations, strains and colour schemes as in Figure 3D. Experiments carried out in triplicate with individual data points, means and standard deviations presented. Transformation ratios calculated and statistical analyses carried out as in Figure 3C. ****, p < 0,001; ***, p < 0,005; **, p < 0,01, n.s., p > 0,05. (C) Competence induction of *radA* is not required for transformation of *rpsL41* point mutation from genomic DNA. tDNA concentrations as in Figure 3D. Strains tested; wildtype (R1501, green), *radA^-^* (R2091, gray), *radA^X-3^* (R4091, orange) and *radA^X-8^* (R4635, purple), *radA^X-3^, CEP_lac_-radA* (R4091, light green) and *radA^X-8^, CEP_lac_-radA* (R4635, light blue). Experiments carried out in triplicate with individual data points, means and standard deviations presented. Transformation ratios calculated and statistical analyses carried out as in Figure 3C. **, p < 0,01; *, p < 0,05, n.s. (not significant), p > 0,05.

### σ^X^ induction of *recA* and *radA* improves tolerance to MMS-mediated genotoxic stress

Next, we explored the importance of transcriptional induction of the *recA* and *radA* operons for genome maintenance by DNA repair. We recently showed that competence provides cells with increased tolerance faced with transient exposure to lethal doses of distinct genotoxic stresses^16^. Although the competence protein ComM, which promotes a transient division delay in competent cells^43^, played a general role in this tolerance, other unknown competence factors were also involved in increasing tolerance to such specific stresses. To determine if HR-mediated DNA repair by RecA and RadA was involved in improving tolerance of competent cells to genotoxic agents, we compared tolerance to lethal doses of MMS and norfloxacin in competent and non-competent cells in either wildtype cells or those lacking *recA* or *radA*, or unable to induce expression of these genes during competence. Competent or non-competent cells were exposed to these genotoxic agents at lethal concentrations^16^ for 60 minutes, before plating to compare colony-forming units (cfu) to determine survival as a percentage. Survival of competent and non-competent populations was then compared to define a tolerance ratio, as previously defined^16^ (Figure 5A).

**Figure 5:**
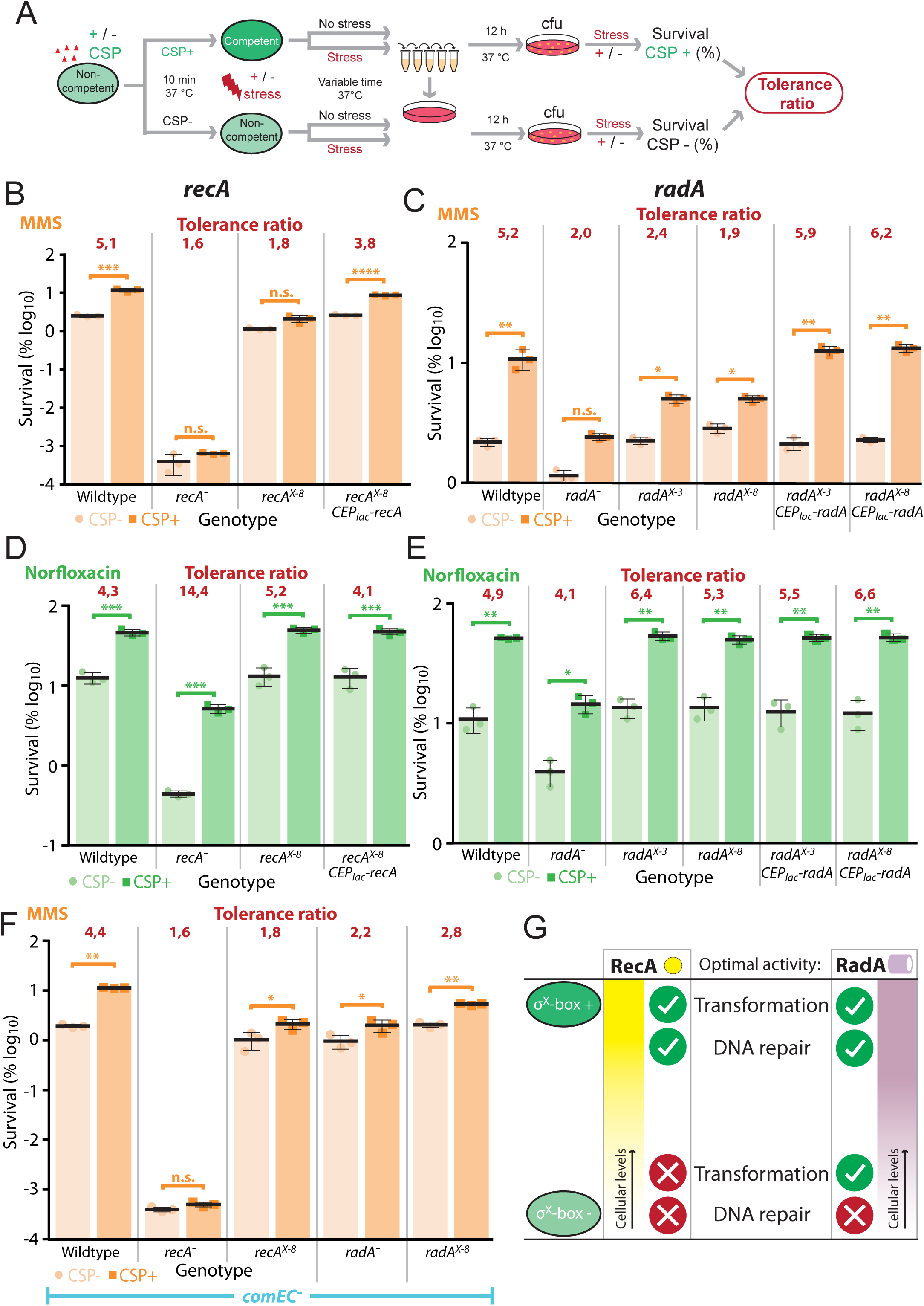
Competence induction of *recA* and *radA* is required for optimal survival of competent cells exposed to MMS. (A) Schematic representation of experimental procedure for survival assay used to gauge the importance of *recA* and *radA* competence induction on tolerance in response to genotoxic stresses^16^. (B) Survival of CSP- (light orange) and CSP+ (dark orange) cells exposed to MMS (625 µg mL^-1^). Wildtype (R1501), *recA^-^* (R4857), *recA^X-8^* (R5077) and *recA^X-8^,* CEP*_lac_-recA* (R5078) strains used. Experiments carried out in triplicate with individual data points, means and standard deviations presented. Asterisks represent significance between transformation efficiencies. ***, p < 0,005. (C) Survival of CSP- (light orange) and CSP+ (dark orange) cells exposed to MMS (625 µg mL^-1^). Wildtype (R1501), *radA^-^* (R2091), *radA^X-3^* (R4091), *radA^X-8^* (R4635), *radA^X-3^,* CEP*_lac_-recA* (R4780), *radA^X-8^,* CEP*_lac_-recA* (R4781) strains used. Representations as in panel B. Asterisks represent significance between transformation efficiencies. *, p < 0,05; **, p < 0,01. (D) Survival of CSP- (light green) and CSP+ (dark green) cells exposed to norfloxacin (100 µg mL^-1^). Strains and representations as in panel B. Asterisks represent significance between transformation efficiencies. n. s., (not significant), p > 0,05; ***, p < 0,005; ****, p < 0,001. (E) Survival of CSP- (light green) and CSP+ (dark green) cells exposed to norfloxacin (100 µg mL^-1^). Strains and representation as in panel C. Asterisks represent significance between transformation efficiencies. n. s., not significant; *, p < 0,05; **, p < 0,01. (F) tDNA acquired from lysed cells is not required for the observed competence mediated increase in tolerance to MMS. Experiments repeated in cells lacking the *comEC* gene, encoding the transformation pore. *comEC^-^* (R2300), *comEC*^-^ *recA^-^* (R5187), *comEC^-^ recA^X-8^* (R5188), *comEC^-^ radA^-^* (R4718) and *comEC^-^ radA^X-3^* (R4720) strains used. Representations as in panel B. Asterisks represent significance between transformation efficiencies. n. s., not significant; *, p < 0,05; ***, p < 0,005. (G) Diagram representing the importance of competence induction of *recA* and *radA* on optimal transformation and DNA repair mechanisms. While competence induction of *recA* is required for both optimal transformation and DNA repair, competence induction of *radA* is required only for optimal DNA repair. Green oblongs represent competent cells with either the *recA* or σ^X^-box intact (σ^X^-box +) or mutated (σ^X^-box -).

MMS causes dsDNA breaks and stalls replication forks by alkylating and depurinating nucleotides in DNA^44,45^, a lethal stress that we recently reported to be better tolerated by competent cells^16^. Compared to wt, the *recA^-^* strain was hyper-sensitive to MMS, and induction of competence did not increase its survival (Figure 5B). This shows that RecA was key for tolerance to MMS in both competent and non-competent cells. Also, while non-competent *recA^X-^*^8^ cells showed similar survival levels to wt cell treated with MMS, the no longer exhibited improved tolerance to MMS upon competence induction (Figure 5B). This indicates that σ^X^ induction of *recA* was required for increased tolerance to MMS. RecA complementation via the ectopic CEP*_lac_-recA* construct recovered the tolerance increase in competent cells (Figure 5B), showing that *recA* alone, rather than any other gene in the same operon (Figure 2A), was responsible for this tolerance increase. The *radA^-^* strain was slightly more sensitive than wt to MMS, but again much less so than *recA^-^*. In addition, the competence-induced tolerance increase was minor in this strain, pointing to an active role for RadA along with RecA in this mechanism (Figure 5C). The *radA^X-3^* and *radA^X-8^* mutant strains also showed tolerance ratios lower than wt (Figure 5C), showing that in the absence of σ^X^ induction of the *radA* operon, competence still increased tolerance of these strains to MMS, but significantly less than wt. Competence induction of the *radA* operon is thus important for improved tolerance to MMS. To explore whether loss of induction of *radA*, rather than any other member of the *radA* operon (Figure 2B), was responsible for this effect, complementation mutants were tested. Indeed, these two strains (*radA^X-3^*, CEP*_lac_-radA*; *radA^X-8^*, CEP*_lac_-radA*) showed tolerance ratios comparable to wt (Figure 5C), confirming that σ^X^-induction of *radA* alone was important for the competence-mediated tolerance to MMS. The competence induction of key HR genes is thus important for the increased tolerance observed in competent cells transiently exposed to lethal concentrations of the genotoxic agent MMS.

To explore whether a similar effect as observed for other DNA-damaging agents, we repeated these tests with norfloxacin. Norfloxacin is an antibiotic which inhibits DNA gyrase and thus prevents DNA replication^46,47^. As shown previously^16^, competent cells survived better than non-competent cells transiently exposed to the antibiotic norfloxacin at lethal concentrations (Figure 5DE). First, the *recA^-^* strain showed significantly reduced survival compared to the wt strain in both competent and non-competent conditions. However, survival remained high above that of the same strain exposed to MMS (Figure 5B). Of note, a marked increase in survival of competent cells was observed in both wt and *recA^-^*strains, another contrast with results observed with MMS. Thus, RecA is important for survival of cells exposed to norfloxacin, due to its involvement in HR-mediated DNA repair, but other RecA-independent pathways also exist to counteract the damaging effect of norfloxacin. In addition, the presence of RecA was not required for the competence-mediated increase in tolerance. Next, we observed that the *recA^X-8^*strain and the complemented *recA^X-8^*, CEP*_lac_-recA* strain both displayed norfloxacin survival profiles and tolerance ratios similar to wt (Figure 5D). This result showed that competence induction of *recA* was not required for improved tolerance to norfloxacin in these conditions. Second, a much smaller decrease in survival following norfloxacin exposure was observed for the *radA*^-^ strain in comparison to the *recA*^-^ strain (Figure 5C), showing that RadA plays a minor role in the repair of norfloxacin-mediated damage. Both *radA^X-3^* and *radA^X-8^* mutants, as well as two complementation strains (*radA^X-3^*, CEP*_lac_-radA*; *radA^X-8^*, CEP*_lac_-radA*) displayed a tolerance profile and ratio similar to wt (Figure 5C), showing that like *recA*, the induction of *radA* during competence was not required for improved tolerance to norfloxacin in these conditions.

Taken together, these results show that the σ^X^-mediated transcriptional induction of the two key HR genes *recA* and *radA* is important for the improved tolerance of competent cells to lethal concentrations of MMS, but not norfloxacin.

### Exogenous DNA internalised by transformation is not required for tolerance to MMS

Since exposure to lethal doses of genotoxic agents kills most cells in a population, competent survivors could use homologous DNA released from neighbouring cells for HR-directed chromosome repair by transformation, and access to this DNA could therefore play a role in the increased tolerance of competent cells to genotoxic stress. Indeed, a previous study suggested that addition of tDNA could improve survival of competent cells faced with the genotoxic stress mitomycin C^15^. However, tDNA can cause agglutination of competent cells, so comparing survival of DNA+ and DNA- conditions using colony-forming units (cfu) can be problematic. To circumvent this issue, we inactivated the *comEC* gene, encoding the transformation pore, rendering cells unable to take up environmental DNA, and repeated the MMS tolerance assay. This allowed us to test the hypothesis of a role for exogenous DNA in tolerance without adding exogenous DNA to the cells, overcoming potential agglutination issues. Results showed that the absence of ComEC did not alter the increased tolerance of competent wildtype cells faced with MMS (Figure 5F). Accordingly, the survival and tolerance ratios of *recA* and *radA* mutants was not affected by the absence of ComEC either. Thus, we conclude that uptake of exogenous DNA by transformation is not necessary for the increased tolerance of competent cells transiently exposed to lethal doses of MMS. This shows that the DNA repair occurring in competent cells surviving lethal MMS exposure uses endogenous chromosomal DNA as a template for HR.

## Discussion

In this study, we explored the competence induction of HR genes *recA* and *radA*, and its importance for their roles in natural transformation and genome maintenance (Figure 5G). We confirmed a previous observation^23^ that competence induction of *recA* was required for optimal transformation efficiency. In contrast, this was not the case for *radA*, where abrogation of competence induction did not significantly alter transformation efficiency, irrespective of the transformation donor or concentration tested. Competence induction of both *recA* and *radA* was required for full competence-induced tolerance to the genotoxic agent MMS. Competence has been historically linked to transformation, but is also associated with many other processes such as biofilm formation^24,25^, virulence^26–29^ and host transmission^30^. Recently, competence was shown to increase tolerance to a wide variety of stresses including genotoxic stress^16^. The finding that competence induction of *recA* and *radA* was important for increased tolerance to genotoxic stress further reveals competence as a key process integrated into the lifestyle of the pneumococcus.

### Competence induction of *recA* and *radA* in transformation

The *recA* and *radA* genes are unusual among competence-induced genes involved in transformation in that they are each controlled by two promoters, one ensuring expression of these genes outside of competence and another, competence-specific promoter (Figure 2AB). This may be because of their roles in both genome maintenance, which occurs in all cells, and transformation, which is restricted to competent cells. We found here that while the competence induction of *recA* was required for optimal transformation, this was not the case for *radA*. This suggests that high cellular levels of RecA are required for transformation, while optimal transformation can occur with very little RadA present. This difference could be explained by the fact that RecA is required for protection of tDNA after internalisation^20^. Preventing the boost of cellular RecA provided by competence could increase the degradation of tDNA by endogenous nucleases and in turn decrease transformation efficiency. Since the role of RecA during HR for transformation is to polymerise on tDNA and search for homology in the recipient chromosome to carry our HR, RecA should polymerise on each tDNA molecule internalised during transformation. In contrast, hexamers of RadA are thought to intervene on D-loop structures, extending them in both directions^22^ after chromosomal integration has occurred. This restricts the role of RadA to HR events where homology search has been successful. Thus, the cellular levels of RadA in competent cells could be less critical to the protection and recombination of tDNA compared to RecA, possibly explaining the observed difference. The fact that ectopic expression of *CEP_lac_-recA* did not fully complement the transformation deficit of the *recA^X-8^* mutation also suggests that high levels of RecA are required to ensure optimal protection of tDNA, since expression levels from the P_lac_ promoter are lower than during competence (Figure S1). In conclusion, the competence induction of *recA* is important for optimal transformation, but this is not the case for *radA*. This is most likely due to a dose effect, with higher levels of RecA required in competent cells due to the nature of the roles of these HR proteins.

### Competence induction of *recA* and *radA* in tolerance to genotoxic stress

The competence induction of *recA* and *radA* was required for optimal tolerance to MMS, but not norfloxacin (Figure 5A-E). This difference may come from the nature of the DNA damages mediated by these two stresses. Norfloxacin inhibits the DNA gyrase, which blocks DNA replication^47^, while MMS alkylates or depurinates nucleotides in DNA, causing base mispairing as well as replication blocks^45^. It may be that survival faced with MMS requires higher cellular levels of HR mediators than in the case of norfloxacin, at least in the conditions tested here. Inactivation of recA rendered cells much more sensitive to MMS than norfloxacin, showing that RecA-mdiated HR is more important for survival of cells exposed to MMS than norfloxacin. In addition, inactivation of *recA* reduced the tolerance ratio of cells faced with MMS, but not norfloxacin, suggesting RecA to be more important for competence-mediated MMS tolerance (Figure 5BC). Removing *recA* or *radA* competence induction did not have the same effect on MMS tolerance. Removing competence induction of *recA* reduced tolerance to non-competent levels (Figure 5B), while both *radA* induction mutants displayed intermediate tolerance levels (Figure 5C). This suggests that similarly to transformation, higher cellular levels of RecA are required for tolerance to MMS compared to RadA. This may be again due to the nature of these HR mediators, with RecA polymerising on ssDNA and RadA promoting branch migration, a role that relies on fewer individual proteins. In addition, while RecA is the central actor of HR, RadA is an accessory protein, and other helicases may be able to compensate for the absence of RadA to facilitate HR, such as RecG or RuvAB^48–50^, which both carry out branch migration during various HR pathways. The uptake of environmental DNA by transformation was not required for competence-mediated tolerance to MMS. This demonstrates that homologous DNA from killed neighbouring cells is not used to repair the genome of competent-tolerant cells by HR. As a result, HR pathways must use endogenous DNA as a template for DNA repair during tolerance to MMS, meaning tolerant cells are self- sufficient for repair of damaged DNA by HR. Interestingly, ectopic expression of *recA* or *radA* in non-competent cells did not alter the survival of these cells, unlike their competence induction (Figure 5). This could be because boosting one of these HR mediators without the other does not increase survival when faced with MMS. Alternatively, the ComM-mediated cell division delay promoted during competence^43^, which is key for increased tolerance to numerous antibiotics and genotoxic agents^16^, could be necessary to give the cells time to reap the beneficial effect of HR-mediated DNA repair. The fact that inactivation of either *recA* (Figure 5B) or *comM*^16^ reduces the tolerance ratio to MMS to almost non-competent levels adds weight to this hypothesis. The transient division delay mediated by ComM improves tolerance to a wide variety of antibiotics and genotoxic agents targeting different cellular processes^16^, and it may be that this delay provides cells with more time to overcome issues caused, with specific mechanisms involved in each case, such as HR-mediated DNA repair in this case.

### Conclusions and perspectives

This study reveals that while the competence induction of the HR gene *recA* is important for transformation, this is not the case for the HR helicase gene *radA*. Pneumococcal competence has historically been linked with transformation, but recently, competence has been shown to provide many more properties to pneumococcal cells. Competence increases tolerance to many stresses, with a ComM-mediated division delay key to this^16^. This study identifies competence induction of *recA* and *radA* genes as important for tolerance of cells faced with transient exposure to lethal doses of MMS. These results further demonstrate that competence provides cells with benefits reaching beyond transformation. Historically, the competence induction of *recA* and *radA* has been assumed to be to ensure their roles in transformation. Here we shown that their induction is at least equally important for DNA repair in competent cells, reinforcing the idea of competence as a general response to cellular stresses, with transformation being only one facet of this response.

## Materials and Methods

### Bacterial strains, competence and transformation

The pneumococcal strains, primers and plasmids used in this study can be found in Table S1. Standard procedures for transformation and growth media were used^51^. In this study, cells were prevented from spontaneously developing competence by deletion of the *comC* gene (*comC0*)^52^, rendering cells unable to produce CSP. Unless described, pre-competent cultures were prepared by growing cells to an OD_550_ of 0.1 in C+Y medium (pH 7) before 10- fold concentration and storage at −80°C as 100 μL aliquots. Antibiotic concentrations (μg mL^−1^) used for the selection of *S. pneumoniae* transformants were: chloramphenicol (Cm), 4.5; erythromycin, 0.05; kanamycin (Kan), 250; spectinomycin (Spc), 100; streptomycin (Sm). Transformation for strain construction was carried out as previously described^51^, with modifications. Briefly, 100 μL aliquots of pre-competent cells were resuspended in 900 μL fresh C+Y medium with 100 ng mL^−1^ CSP (and 50 µM IPTG where necessary), then incubated at 37°C for 10 min. tDNA was then added to a 100 μL aliquot of this culture, followed by incubation at 30°C for 20 min. Cells were then diluted and plated on 10 mL CAT agar with 5% horse blood before incubation at 37°C for 2 h. A second 10 mL layer of CAT agar with appropriate antibiotic was added to plates to select transformants, and plates without antibiotic were included for comparison to calculate transformation efficiency. Plates were incubated overnight at 37°C. For the generation of point mutants, after exposure to DNA, cells were rediluted in 1,4 mL C+Y medium and incubated at 37 °C for 3h 30 min, before dilution and plating without selection. Successful transformants were identified by PCR and sequencing. Strains possessing the P*_lac_* promoter platform were grown in 50 µM IPTG from the beginning of growth, as previously described^26^. GraphPad Prism was used for statistical analyses. To generate strain R2191, strain R1502 was transformed with genomic DNA from strain *ΔradA*^19^, with *radA::spc* transformants selected using spectinomycin. To generate strain R2300, R1501 was transformed with a mariner mutagenesis fragment of the *comEC* gene, generated using primer pair CJ974-CJ975, with the pR410 plasmid, as previously described^53^. A *comEC::kan* transformant with a kanamycin resistance cassette inserted in the *comEC* gene in a co- transcribed orientation was isolated by selection with kanamycin. To generate strain R4091, PCR fragments of regions upstream and downstream of the *radA* operon σ^X^ box were amplified, with 3 bp of the σ^X^ box mutated using primer pairs MB180-MB183 and MB181- MB182. Splicing overlap extension (SOE) PCR with these two fragments as templates and primer pair MB183-MB182 was carried out to generate a PCR fragment with the *radA* operon σ^X^ box mutated and ∼2 kb of homologous sequence on either side. This DNA fragment was transformed into strain R1818 without selection and transformants were identified by sequencing. To generate strain R4635, PCR fragments of regions upstream and downstream of the *radA* operon σ^X^ box were amplified, with 8 bp of the σ^X^ box replaced with a *Bam*HI site (GGATCC) flanked by two other bases using primer pairs DDL122-DDL123 and DDL124- DDL125. SOE PCR with these two fragments as templates using primer par DDL122-DDL125 was carried out to generate a PCR fragment with the *radA* operon σ^X^ box mutated and ∼2 kb of homologous sequence on either side. This DNA fragment was transformed into strain R1501 without selection and successful transformants were identified by amplification of the region using primer pair DDL122-DDL125 followed by restriction by *Bam*HI for 1 h at 37 °C. To generate strain R4660, the *radA* gene was amplified from R1501 using primer pair oALS- CJ684 possessing a *Bsp*HI site upstream and a *Bgl*II site downstream, for ligation with *Nco*I and *Bam*HI respectively, since *radA* possessed both of those sites in its sequence. Upstream *CEP_lac_* sequence flanked by *Sal*I and *Nco*I sites was amplified from R3833 using primer pair CJ588-CJ680. These two DNA fragments were ligated into the pCEP*R*-*luc* plasmid^42^, digested with *Sal*I and *Bam*HI, and the ligation product was transformed directly into R1501, with transformants selected with kanamycin. To generate strain R4664, the same strategy as for strain R4660 was employed, with the *radA* gene replaced with *recA*, amplified from R1501 with primer pair CJ681-CJ682, flanking *recA* with *Nco*I and *Bam*HI restriction sites. To generate strain R4780, strain R4091 was transformed with gDNA from strain R4660, and transformants selected with kanamycin. To generate strain R4781, strain R4635 was transformed with gDNA from strain R4660, and transformants selected with kanamycin. To generate strain R5077, the *recA^X-8^* mutation (the *recA* promoter σ^X^ box replaced with AGGTACCT, also termed *cinA^cinbox-^* ^40^) was amplified from strain R4446 using primer pair DDL105-DDL108 and transformed into R1501 without selection. Successful transformants were identified by amplification of the region using primer pair DDL105-DDL108 followed by restriction by *Bam*HI for 1 h at 37 °C. To generate strain R5078, strain R5077 was transformed with a PCR fragment amplified from R4664 using primer pair CJ574-CJ575 and containing CEP*_lac_-recA*, with transformants selected with kanamycin. To generate strain R5079, strain R4664 was transformed with a PCR fragment amplified from strain R4857 using primer pair CJ808-CJ829 and containing *recA*::*trim*, with transformats selected with trimethoprim.

### Western blot

A time-course Western blot of R1501 (*comC0*) was carried out to explore the induction of *recA* and *radA* during competence. Cells were diluted 100-fold in 10 mL C+Y medium and grown to OD 0.1. A 1 mL sample was taken at t = 0, and competence was then induced by addition of CSP (100 ng mL^-1^). 1 mL samples were then taken 5, 10, 15, 20, 30, 60, 80 and 100 minutes after competence induction, with growth also measured by OD_492_ reading. 1 mL samples were centrifuged (5 min, 5,000 g) and pellets were resuspended in 40 µL of TE 1 x supplemented with 0.01% sodium deoxycholate (DOC) and 0.02% sodium dodecyl sulphate (SDS). Samples were then incubated for 10 minutes at 37°C before addition of 40 µL 2x sample buffer with 10% β-mercaptoethanol, followed by incubation at 85°C for 10 minutes. Samples were then adjusted to normalize the number of cells per sample using the OD_492_ readings, and loaded onto SDS-PAGE gels (BIORAD), migrated for 30 min at 200V, and transferred onto nitrocellulose membrane using a Transblot Turbo (BIORAD). Membranes were blocked for 1h at room temperature in 1x TBS with 0.1% Tween20 and 10% milk, before two washes in 1x TBS with 0.1% Tween20. Membranes were then separated in two at the 25 kDa mark and probed with primary antibodies (1/10,000 dilution of α-RadA or α-RecA antibodies for top membrane half, 1/10,000 dilution of α-SsbB antibodies for bottom membrane half to detect both SsbB and SsbA^38^) in 1x TBS with 0.1% Tween20 and 5% milk overnight at 4°C. After a further four washes in 1x TBS with 0.1% Tween20, membranes were probed with anti-rabbit secondary antibody (1/10,000) for 1h 30 min, followed by another four washes in 1x TBS with 0.1% Tween20. Membranes were activated using Clarity Max ECL (BIORAD) and visualized in a ChemiDoc Touch (BIORAD). To compare the expression profiles of the *recA^X-8^* strain with wildtype and *recA^-^* strains, R1501 (*comC0*), R4857 (*comC0*, *recA^-^*) and R5077 (*comC0*, *recA^X-8^*) cells were diluted 100-fold in 3 mL C+Y medium and grown to OD 0.1. Samples were split into two 1,5 mL volumes (CSP+ and CSP-) and competence was induced in CSP+ tubes by addition of CSP (100 ng mL^-1^). After 15 min at 37°C, samples were centrifuged (5 min, 5,000 g) and then treated as described above. To compare the expression profiles of *radA^X-3^* and *radA^X-8^* strains with wildtype and *radA^-^* strains, R1501 (*comC0*), R2191 (*comC0, radA::spc*), R4091 (*comC0, radA^X-3^*) and R4635 (*comC0, radA^X-8^*) cells were treated in the same manner. To validate the complementation of *recA* competence induction by *P_lac_-recA*, R1501 (*comC0*), R4857 (*comC0, recA::trim*), R5077 (*comC0, recA^X-8^*) and R5078 (*comC0, recA^X-8^, P_lac_-recA*) strains were diluted 100-fold in 3 mL C+Y medium (possessing or not 50 µM IPTG) and grown to OD 0.1. Samples were split into two 1,5 mL volumes (CSP+ and CSP-) and competence was induced in CSP+ tubes by addition of CSP (100 ng mL^-1^). After 15 min at 37°C, samples were centrifuged (5 min, 5,000 g) and then treated as described above. To validate the complementation of *radA* competence induction by *P_lac_-radA*, R1501 (*comC0*), R2191 (*comC0, radA::spc*), R4780 (*comC0, radA^X-3^, P_lac_-radA*) and R4781 (*comC0, radA^X-8^, P_lac_-radA*) strains were diluted 100-fold in 3 mL C+Y medium (possessing or not 50 µM IPTG as shown in Figure 6A) and grown to OD 0.1. Samples were split into two 1,5 mL volumes (CSP+ and CSP-) and competence was induced in CSP+ tubes by addition of CSP (100 ng mL^-1^). After 15 min at 37°C, samples were centrifuged (5 min, 5,000 g) and then treated as described above.

### Transformation efficiency test

To test the efficiency of transformation in wildtype and mutant strains, 100 µL pre-competent cultures were resuspended in 900 µL fresh C+Y medium pH 7.6, and 100 ng mL^-1^ CSP was added. Cells were incubated for 10 min at 37°C, before 100 µL was added to tubes containing desired tDNA. tDNA identity was either a 3,434 bp *rpsL41c* fragment^22^ (Figure 3A) possessing a centrally localized point mutation conferring streptomycin resistance (amplified from R304 with primer pair MB117-MB120), a 4,187 bp *rpsL41e* fragment^22^ (Figure 3A) possessing an eccentric point mutation conferring streptomycin resistance (amplified from R304 with primer pair MB121-MB132) or a 5,000 bp fragment (Figure 4A) possessing a Kan^R^ heterologous cassette (1,340 bp) inserted into the *comD* gene (amplified from R1745 with primer pair CJ402-CJ409). Transforming DNA was added at varying concentrations as indicated ranging from saturating (2,500 pg mL^-1^) to very low (2,5 pg mL^-1^) concentrations. Transforming cultures were incubated at 30°C for 20 minutes, before dilution and plating of appropriate dilutions for selected and non-selected cells in 10 mL CAT agar medium with 3% horse blood. Plates were incubated 2 h at 37°C, before addition of a second 10 mL layer of CAT agar medium possessing either streptomycin (for *rpsL41C* and *rpsL41e* transformants) or kanamycin (for *comD::kan* transformants). Plates were incubated overnight at 37°C and colonies present on selected and non-selected plates were compared to calculate transformation efficiencies in each condition tested.

### Survival assays

To explore the effect of *recA* and *radA* competence induction on tolerance to genotoxic agents, survival assays were carried out as described previously^16^, as shown in Figure 5A. For the *recA* experiments, strains tested were R1501 (*comC0*), R4857 (*comC0, recA::trim*), R5077 (*comC0, recA^X-8^*), R5078 (*comC0, recA^X-8^,* CEP*_lac_-radA*). For the *radA* experiments, strains tested were R1501 (*comC0*), R2191 (*comC0, radA::spc*), R4091 (*comC0, radA^X-3^*), R4635 (*comC0, radA^X-8^*), R4780 (*comC0, hexA::ery, radA^X-3^,* CEP*_lac_-radA*), R4781 (*comC0, radA^X-8^,* CEP*_lac_-radA*). Stresses tested at levels above the minimum inhibitory concentration (MIC)^16^ were norfloxacin (100 µg mL^-1^) and MMS (625 µg mL^-1^). Pre-cultures were diluted 50-fold in 3 mL C+Y medium pH 7 and incubated at 37°C until OD_550_ 0,3. These cultures were rediluted 50-fold in 5 mL fresh C+Y medium pH 7, and incubated at 37°C until OD_550_ 0,1. Cultures were then diluted to OD_550_ 0,004 and split into two 2,5 mL samples, with competence induced in one sample by addition of 100 ng mL^-1^ CSP. Samples were then incubated at 37°C for 15 min to allow competence development, before each being split into two 1 mL samples either exposed to the genotoxic stress or not. A further incubation of 1 h at 37°C was carried out, followed by dilution and plating of cells in 10 mL CAT agar medium with 3% horse blood. Plates were incubated at 37°C overnight, and survival percentages were determined by comparing cfu of stressed and non-stressed cells. Tolerance ratios were calculated by comparing survival of CSP+ and CSP- cells.

## Acknowledgements

This work was funded by the Centre National de la Recherche Scientifique, University Paul Sabatier and the Agence Nationale de la Recherche (grants ANR-10-BLAN-1331 and ANR- 17-CE13-0031).

---

**Figure S1:**
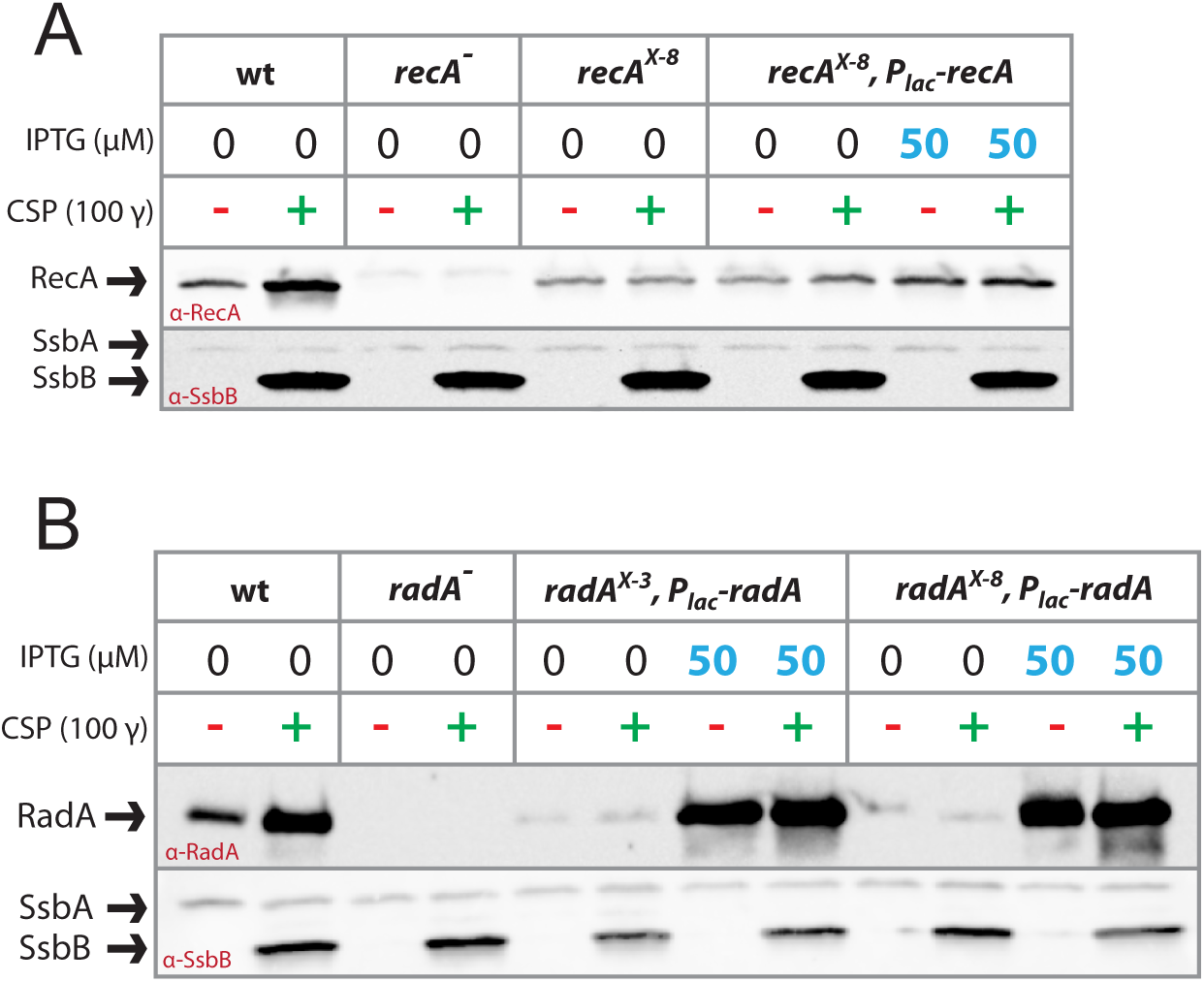
Complementation of *recA* and *radA* competence induction mutants. (A) Complementation of *recA^X-8^* strain by CEP*_lac_-recA*, visualised by Western blot. Wildtype (R1501), *recA^-^* (R4857), *recA^X-8^* (R5077), *recA^X-8^,* CEP*_lac_-recA* (R5078) strains used. α–RecA antibodies used. (B) Complementation of *radA^X-3^* and *radA^X-8^* strains by CEP*_lac_-radA*, visualised by Western blot. Wildtype (R1501), *radA^-^* (R2091), *radA^X-3^,* CEP*_lac_-recA* (R4780), *radA^X-8^,* CEP*_lac_-recA* (R4781) strains used. α–RadA antibodies used.

**Table S1:**
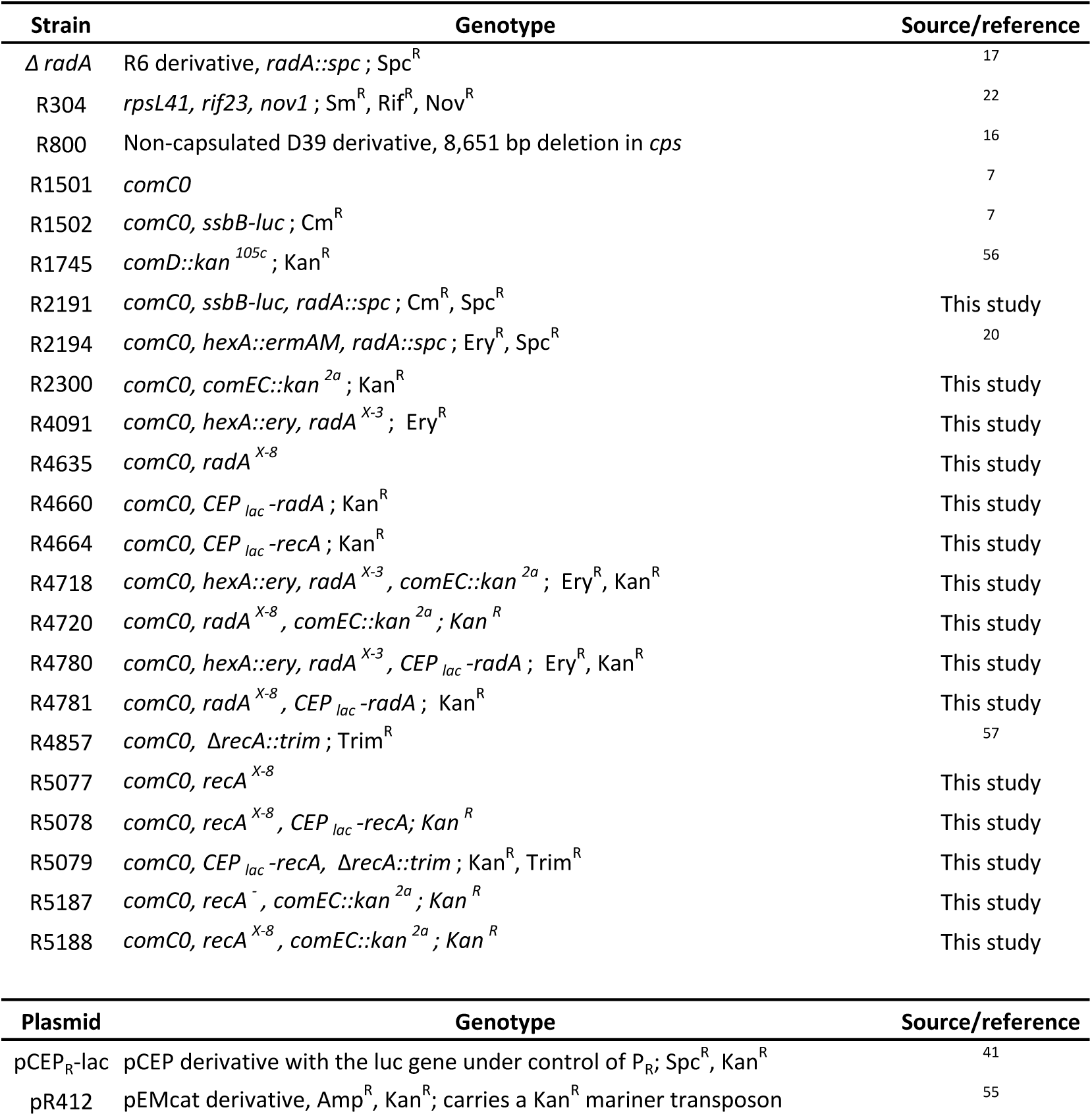
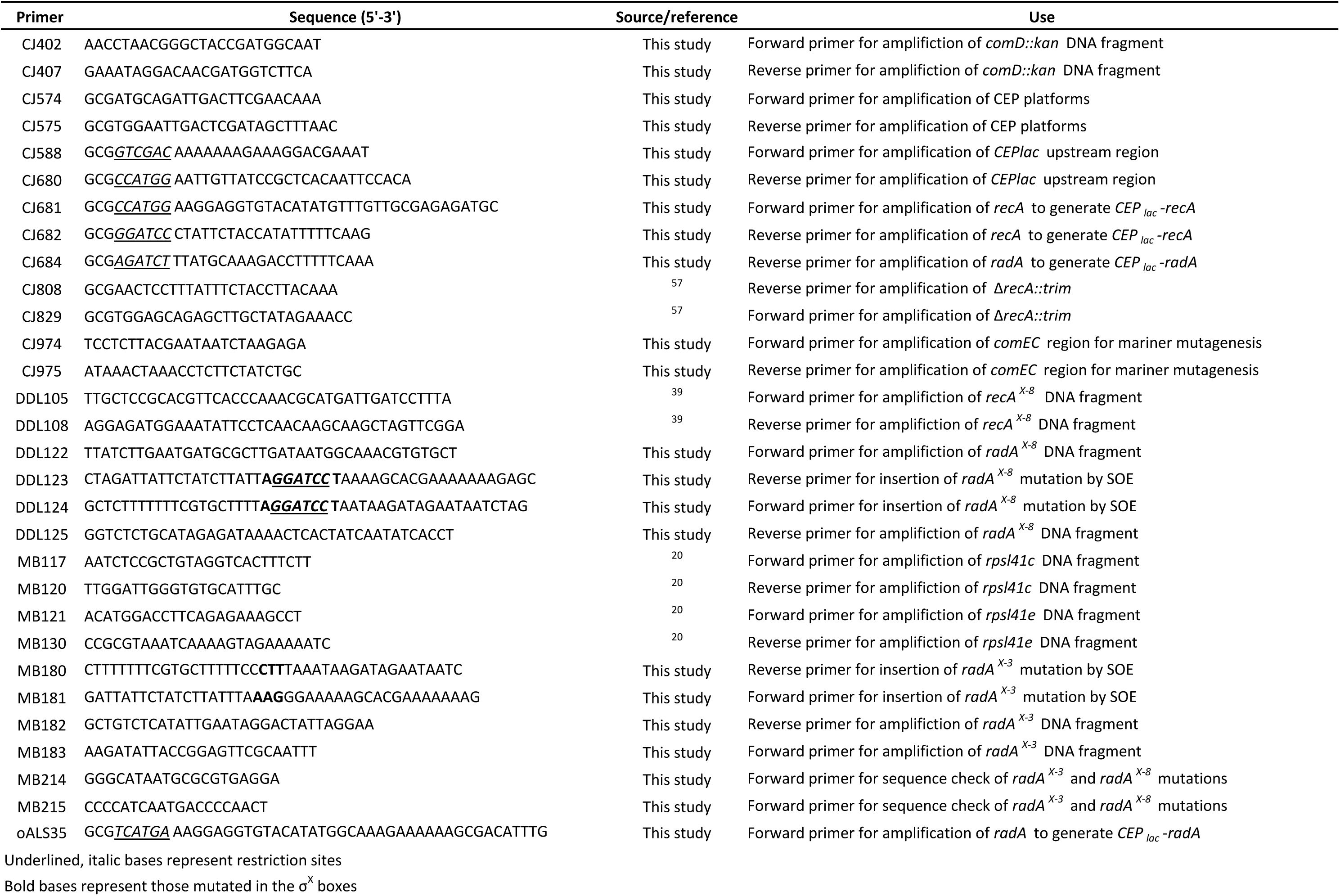
Strains, plasmids and primers used in this study.

